# The cryptic impacts of invasion: Functional homogenization of tropical ant communities by invasive fire ants

**DOI:** 10.1101/611749

**Authors:** Mark K. L. Wong, Benoit Guénard, Owen T. Lewis

## Abstract

Invasive insects represent major threats to ecosystems worldwide. Yet their effects on the functional dimension of biodiversity, measured as the diversity and distribution of traits, are overlooked. Such measures often determine the resilience of ecological communities and the ecosystem processes they modulate. The fire ant *Solenopsis invicta* is a highly problematic invasive species occurring on five continents. Its impacts on the taxonomic diversity of native ant communities have been studied but its impacts on their functional diversity are unknown. Comparing invaded and uninvaded plots in tropical grasslands of Hong Kong, we investigated how the presence of *S. invicta* affects the diversity and distribution of ant species and traits within and across communities, the functional identities of communities, and functionally unique species. We calculated the functional diversity of individual species, including the trait variation from intraspecific polymorphisms, and scaled up these values to calculate functional diversity at the community level. Invasion had only limited effects on species richness and functional richness, which were 13% and 8.5% lower in invaded communities respectively. In contrast, invasion had pronounced effects on taxonomic and functional composition due to turnover in species and trait values. Furthermore, invaded communities were functionally more homogeneous, displaying 23% less turnover and 56% more redundancy than uninvaded communities, as well as greater clustering and lower divergence in trait values. Invaded communities had fewer functionally-unique individuals and were characterized by ant species with narrower heads and bodies and shorter mandibles. Our results suggest that studies based only on taxonomic measures of diversity or indices describing trait variety risk underestimating the full ramifications of invasions. Investigating the diversity and distributions of traits at species, community and landscape levels can reveal the cryptic impacts of alien species which, despite causing little taxonomic change, may substantially modify the structure and functioning of ecological communities.

## Introduction

The seemingly limitless exchange of alien species worldwide (Seebens et al., 2017) is a dominant phenomenon of global change with major implications for both nature and human wellbeing. Invasions by alien species are not only the second most common cause of extinctions (Bellard, Cassey & Blackburn, 2016) but also drive cascading impacts on ecosystems, cause economic damage and undermine human health (Pyšek & Richardson, 2010). As with many ecological phenomena, invasions have long been studied by summarising ecological communities using metrics based on species’ taxonomic identities: abundance, richness and diversity. Changes in these metrics, however, may provide only limited insight into the specific mechanisms (such as novel niches and enemy escape) underlying the causes of invasions (MacDougall, Gilbert & Levine, 2009). Furthermore, the effects of biodiversity on ecosystem processes often depend on the functions performed by the different species, rather than on species numbers and identities *per se* (Gagic et al., 2015). Summarising biodiversity in terms of the traits that directly impact organisms’ ecological interactions (Functional Diversity) (McGill et al., 2006) may advance understanding of the causes and consequences of invasions.

Traits are the specific phenotypic properties of organisms which modulate their ecological interactions. The interactions of key ‘functional traits’ influence organism fitness and may also contribute to ecosystem functions (Wong, Guénard & Lewis, 2018). Thus, studying the diversity of functional traits in an ecological community can simultaneously reveal how biodiversity and ecosystem processes are impacted by disturbances such as invasive species. For instance, trait-based studies of plant and vertebrate communities undergoing invasion by alien species revealed declines in functional richness (the variety of trait values in individual communities) and a tendency for functional homogenization (i.e., an increased similarity in trait values between communities) (Villéger, Grenouillet & Brosse, 2014; Castro-Díez et al., 2016). Crucially, invasion-driven changes in functional structure were linked to altered ecosystem functions (Castro-Díez et al., 2016).

One general limitation of earlier trait-based studies, including those on invasions, has been the exclusive use of mean trait values for species to estimate community functional diversity. This approach underestimates intraspecific trait variation, which can strongly influence community dynamics and ecosystem processes (Des Roches et al., 2018). Intraspecific trait variation has also been implicated in the success of some invasive species (González-Suárez, Bacher, & Jeschke, 2015). Recently-developed statistical tools such as the Trait Probability Density (TPD) framework can incorporate intraspecific trait variation into estimates of functional diversity (Carmona et al., 2016), but few studies have explored this within the context of invasions. Additionally, although trait-based studies are advancing understanding of invasions by plants and vertebrates, there is a shortage of similar work on invasive insect species (Wong et al., 2018), despite their ubiquity and widespread impacts on biodiversity and ecosystem services (Bradshaw et al., 2016).

Many ants, for instance, are contenders for the world’s most harmful invasive species (Lowe et al., 2000). Numerous studies have documented the consequences of ant invasions for native ant communities, which include declines in species richness, taxonomic homogenization, and phylogenetic clustering (reviewed in Lessard et al., 2009). These studies focus on taxonomic and phylogenetic patterns, and rarely consider trait values directly. Few such studies document the phenotypes that were lost or gained at the community level, and none directly assessed the impacts of invasions on the functional diversity of individual communities (functional alpha diversity) or patterns across multiple communities (functional beta diversity). Furthermore, no trait-based studies on ants, and few for insects in general include intraspecific trait variation in estimates of community functional diversity (Wong et al., 2018). Intraspecific trait variation is expected to be high in polymorphic species and it may influence how such species respond to or effect ecological change. Some ant species display marked variation in the body size and morphology of their worker caste (worker polymorphism), a feature which may contribute to colony fitness and ecological success (Tschinkel, 1988; Wilson, 2003). It has been observed that polymorphic ant species can surpass monomorphic ones in their abilities to collect resources varying in size, and to access environments varying in rugosity (Farji-Brener, Barrantes & Ruggiero, 2004); that is, polymorphic species may access a wider variety of niches than monomorphic species. Thus, a basic yet apparently untested assumption is that polymorphic species have higher functional (trait) richness than monomorphic species.

The Red Imported Fire Ant, *Solenopsis invicta* Buren, 1972, is native to South America but its expanding global range already encompasses five continents (Ascunce et al., 2011; Guénard et al., 2017). Furthermore, models indicate that vast areas throughout the tropics and subtropics are susceptible to future invasion (Morrison et al., 2004). *Solenopsis invicta* has been an extensive ecological problem throughout the United States for decades due to its strong impacts on native biodiversity at multiple trophic levels (including native ants) and the subsequent cascading effects on ecosystems (Porter & Savignano, 1990; Vinson, 1997). The species is the second most costly invasive insect in the United States, with economic costs exceeding US$7 billion annually (Bradshaw et al., 2016). *Solenopsis invicta* are dietary generalists and mature colonies consist of a polymorphic worker caste (Tschinkel, 2006), a factor which, alongside their strong interspecific aggression, may contribute to the success of invasive populations (Tschinkel, 1988; 2006). The first report of *Solenopsis invicta* in Asia originated from Taiwan in 2003; the species was later detected in continental China in 2005 (Ascunce et al., 2011). In spite of their devastating impacts elsewhere, studies on the ecology of *S. invicta* invasions in Asia have been limited. Preliminary observations from agrosystems in China suggest *S. invicta* invasions may be associated with declines in the species richness of arthropod communities (Wang et al., 2018).

Here, we investigate the impacts of the invasion of *S. invicta* on taxonomic and functional diversity within (alpha diversity) and between (beta diversity) native ant communities in Hong Kong. This is the first comprehensive trait-based study of an ant invasion’s impact on the functional facet of biodiversity, as well as the first to incorporate polymorphisms in calculations of functional diversity at the species level.

At the scale of the individual community, we examined how invasion by *S. invicta* affected (i) species and functional richness, (ii) abundance-weighted indices of multidimensional functional diversity, and (iii) functional identity, the dominant value of a trait in the community. We predicted lower species and functional richness as well as altered functional identities in communities invaded by *S. invicta* (Porter & Savignano, 1990; Castro-Díez et al., 2016). At the multi-community (landscape) scale, we investigated how *S. invicta* invasion affected taxonomic and functional beta diversity (the dissimilarities in species and traits between communities). We predicted that invasion by *S. invicta* would lead to taxonomic homogenization (Lessard et al., 2009), which would be associated with functional homogenization, as observed for other alien taxa (Villéger et al., 2014). Using measures of species-level functional diversity, we also identified functionally unique species and compared their relative uniqueness to uninvaded and invaded communities. Lastly, we measured and compared the functional richness of different species, with the prediction that the functional richness of polymorphic species such as *S. invicta* would exceed that of monomorphic species.

## Materials and methods

### Study area and sampling design

Our study sites are two (<2 km apart) wetland reserves in northern Hong Kong: Lok Ma Chau (22.512°N, 114.063°E) and Mai Po (22.485°N, 114.036°E). Both reserves encompass abandoned fish farms that have since been conserved for >35 years as habitats for migratory birds. Each contains a network of bunds (width ≤5 m) which separate individual ponds (Fig. S1). The habitat is relatively homogeneous and comprises exposed grasslands with native tree species interspersed throughout. Ant communities in this landscape are comprised mostly of native species but pilot surveys from 2015 to 2017 revealed that colonies of *S. invicta* are present at high densities at multiple locations. We marked these locations, and in 2018 selected a total of 61 plots, each a 4 × 4 m quadrat, to reflect two ant community types: communities with *S. invicta* absent (uninvaded; 37 plots), and those with *S. invicta* present (invaded; 24 plots). A minimum distance of 20 m between individual plots facilitated independent observations since most ant species in the region forage no further than 5 m from their nests (Eguchi, Bui & Yamane, 2004) and *S. invicta* forage within 4 m of their nests (Weeks, Wilson & Vinson, 2004). Given the homogeneity of the landscape we assumed that any community differences observed between uninvaded and invaded plots would primarily be a consequence of invasion by *S. invicta*; environmental data collected at fine spatial resolutions were used to test this assumption (see below). Sampling was conducted from April to September 2018. At each plot, six pitfall traps (55 mm in diameter) were installed to sample the ant community over 48 hours. All specimens were sorted into morphospecies and subsequently most were identified to species using taxonomic keys.

### Environmental data

We used local GIS models (Morgan & Guénard, 2019) to obtain high-resolution data (30 × 30 m rasters) for three environmental variables corresponding to each plot: Normalized Difference Vegetation Index (NDVI), mean annual temperature, and mean annual precipitation.

### Assembling the individual-level trait dataset

Here we aimed to obtain values of functional diversity that incorporated intraspecific trait variation, including the variation arising from worker polymorphisms. We assembled an individual-level trait dataset comprising data for seven morphological traits that regulate ant physiology and behaviour and that are hypothesized to impact performance and fitness (Table 1). Using mounted specimens from the pitfall traps and a Leica M165c stereo microscope paired with Leica Application Suite software, we recorded high-resolution images and performed trait measurements on at least 10 individual workers of every species (N=319). For dimorphic species of *Camponotus* and *Pheidole* where workers comprise two distinct sub-castes (minors and majors), we included trait data for individuals of both sub-castes based on the relative proportions (i.e., ratio of minors to majors) observed in natural colonies (Passera, 1984; Wilson, 2003). The invader *S. invicta* has a polymorphic worker caste, and Tschinkel (1988) showed that this polymorphism is mainly expressed in the morphological variation displayed by the ‘majors’ (head width >0.7 mm), which are present only in mature colonies where they comprise 35% of the worker population; ‘minors’ of head width <0.7 mm comprise the remaining 65% in mature colonies and juvenile colonies only consist of minors. We observed that all invaded plots contained majors (head width >0.7 mm), suggesting they were mature colonies; thus, our trait data for *S. invicta* (n=20) included both minors (65% of individuals) and majors (35% of individuals).

**Table 1.** The seven traits measured on each individual, and each trait’s hypothesized links to the performance and fitness of ants.

| Trait | Measurement | Hypothesized link to performance and fitness |
| --- | --- | --- |
| *size* | Weber's length:<br>diagonal length of<br>mesosoma | Modulates vital and physiological rates,<br>determines physical constraints and exposure to<br>predators, influences resource type and<br>acquisition efficiency (Silva & Brandão, 2010). |
| *head width* | Width of head<br>including eyes | Determines the size of gaps through which an<br>individual can pass (Schofield, Bishop & Parr,<br>2016) and the volume of muscles powering the<br>mandibles during foraging (Richter et al., 2019). |
| *eye width* | Width of left eye | Determines ability in navigation, foraging,<br>predator and prey detection, and indicative of<br>activity times (Silva & Brandão, 2010). |
| *mandible length* | Length of left<br>mandible | Responds to selection on diet type and<br>specialization (Silva & Brandão, 2010). |
| *scape length* | Length of scape of<br>left antenna | Responds to selection on navigation and sensory<br>abilities (Silva & Brandão, 2010). |
| *pronotum width* | Width of pronotum | Determines volume of muscles for head-support<br>and load-bearing (Keller et al., 2014). |
| *leg length* | Combined length of<br>femur and tibia of left<br>hind leg | Determines mobility; leg length influences<br>running speed, which affects success in foraging<br>or escape from predators (Grevé et al., 2019). |

### Compressing trait variation

We divided the measurements of six traits (*head width*, *eye width*, *mandible length*, *scape length*, *pronotum width* and *leg length*) by *size* to reduce correlation with body size. We then log-transformed the values of all seven traits to reduce the influence of extreme values, and standardized trait values to have mean of zero and unit variance. Next, we used Principal Components Analysis (PCA) to synthesize the major axes of variation in multidimensional trait space and to reduce the number of dimensions used to calculate functional diversity indices. We performed the PCA using the mean trait values of each species and subsequently predicted the values of the PCA components for all individuals in the dataset. We used species means instead of individual trait values in the PCA because using the latter could bias the analysis if some species had disproportionately large numbers of individuals in the dataset. We retained the first two components of the PCA, which had eigenvalues greater than unity, and which captured 76.9% of the total variance in traits. We then predicted the values of these two components for every individual in the trait dataset and used these new ‘traits’ to calculate functional diversity indices.

### Functional diversity from species to communities

All functional diversity indices were calculated using the Trait Probability Density framework which incorporates intraspecific variation, the multidimensional nature of traits, species abundances, and probabilistic trait distributions (see Carmona et al., 2016). First, we used multidimensional probability density functions to calculate trait probability distributions (which reflect the probabilities of observing different trait values) at the level of individual species (TPDsp). Next, we scaled up TPDsp to local community levels (TPDcom) by summing the TPDsp of all species in each local ant community, weighted by their relative abundances – which we estimated as frequencies of occurrence. Finally, five different indices for functional diversity were calculated using each community’s TPDcom. The indices were Functional Richness (FRic), the volume of functional space occupied by the community; Functional Evenness (FEve), the regularity of the distribution of abundance in functional space; Functional Divergence (FDiv), a measure of how abundances tend to be on the outer margins of the functional space while controlling for functional richness; Rao, the abundance-weighted dispersion of individuals (or species) in functional space; and Functional Redundancy (FRed), the degree to which trait values are represented by multiple species in the community (Carmona et al., 2016). We used this multi-index approach to measure functional diversity because no one index can encapsulate the independent components of functional diversity (Mouchet et al., 2010). In addition to calculating the observed values of the functional diversity indices, we calculated Standardized Effect Size (SES) values for all indices so as to estimate community-level functional diversity that had been corrected for potential effects of species richness (Swenson, 2014). SES values were calculated by comparing the observed values to values generated from 999 constrained null models randomizing the community data matrix using the “Independent Swap” algorithm. The formula for calculating SES is:

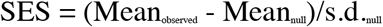

### Functional identity

To estimate functional identity, we calculated the community-weighted mean (CWM) for each trait in every local ant community. A CWM reflects the dominant value of a given trait in a given community (Swenson, 2014). We calculated CWMs using mean trait values of species weighted according to their relative abundances in the different communities. Size-correction was applied to all traits except *size* (see above).

### Taxonomic and functional beta diversity

We calculated six pairwise measures of taxonomic and functional beta diversity for all possible pairs of local ant communities. We used matrices of species’ abundances to calculate the Taxonomic Dissimilarity between each pair, and further decomposed this into Taxonomic Turnover (dissimilarities arising from the replacement of species between communities), and Taxonomic Nestedness (dissimilarities in the relative abundances of species that occurred in both communities). We calculated the Functional Dissimilarity between paired communities using their TPDcom, and further decomposed this into Functional Turnover (dissimilarities in the trait values between communities), and Functional Nestedness (dissimilarities in the relative abundances of trait values shared between communities). In addition to observed values, we calculated SES values (from comparisons with 999 constrained null models using the “Independent Swap” algorithm) for all components of functional beta diversity. Although the “Independent Swap” algorithm may not be optimal for generating null models of beta diversity patterns shaped by dispersal limitation (Swenson, 2014), this is unlikely to be a problem for the present study because all species disperse by flying alates that can travel distances exceeding the scale of the study landscape (the maximum distance between any two plots was 4 km).

### Species’ functional richness and functional uniqueness

We calculated functional richness and functional uniqueness values for all species. Functional richness was calculated based on each species’ trait probability distribution (TPDsp) (Carmona et al., 2016). Species’ functional uniqueness values were calculated relative to individual local communities, based on the degree to which a species’ functional space (TPDsp) did not overlap with a local community’s functional space (TPDcom) (Carmona et al., 2016). We calculated each species’ ‘relative uniqueness’ with respect to the different uninvaded and invaded communities, as well as its ‘objective uniqueness’ in the species pool (using a hypothetical community containing all species at equal abundance).

### Statistical analysis

#### Taxonomic alpha diversity, functional diversity and CWMs

We used separate linear mixed-effects models to assess whether the values of alpha taxonomic diversity, functional diversity indices (including observed and SES values) and CWMs differed significantly between uninvaded and invaded local ant communities, while including a random effect of environmental variation that was captured in the first component of a PCA for the three environmental variables, which had eigenvalues greater than unity.

#### Taxonomic and functional beta diversity

We used non-metric multidimensional scaling (NMDS) to scrutinize the relationships between and among invaded and uninvaded local ant communities in multidimensional space (Fig. S2). We used PERMANOVA (9,999 permutations) to quantify dissimilarity, turnover and nestedness between the observed taxonomic and functional compositions of uninvaded and invaded communities. We used permutation tests for multivariate dispersions to assess whether the levels of taxonomic and functional beta diversity (in three components) observed among uninvaded communities differed from those observed among invaded communities. We also used nonparametric Mann-Whitney U tests to compare SES values of the three functional beta diversity components between uninvaded and invaded communities.

#### Functional uniqueness of individual species

We calculated each species’ average relative uniqueness to uninvaded and invaded communities and regressed these against its objective uniqueness in a linear model

#### Software

We used the following packages in R software version 3.3.3 (R Core Team, 2017): *TPD* (Carmona, 2018) for calculating trait probability distributions, functional diversity indices, functional dissimilarity and functional uniqueness measures, *FD* (Laliberté, Legendre & Shipley, 2014) for calculating CWMs, *betapart* (Baselga et al., 2018) for beta diversity analyses, *lme4* (Bates et al., 2015) for linear mixed-effects models, *MASS* (Venables & Ripley, 2002) for NMDS, and *ggplot2* (Wickham, 2009) for the production of graphics.

## Results

### Community composition and species richness

A total of 29 ant species (including *S. invicta*) were collected from 366 pitfall traps in 37 uninvaded plots and 24 invaded plots (Table S1). The species composition across invaded and uninvaded communities was similar overall, with 27 of the 28 native species occurring in both community types, and only one species not found in invaded communities. On average, the species richness of invaded communities was marginally and non-significantly lower (by 13%) than that of uninvaded communities (Table 2; Fig. 1).

**Table 2.**
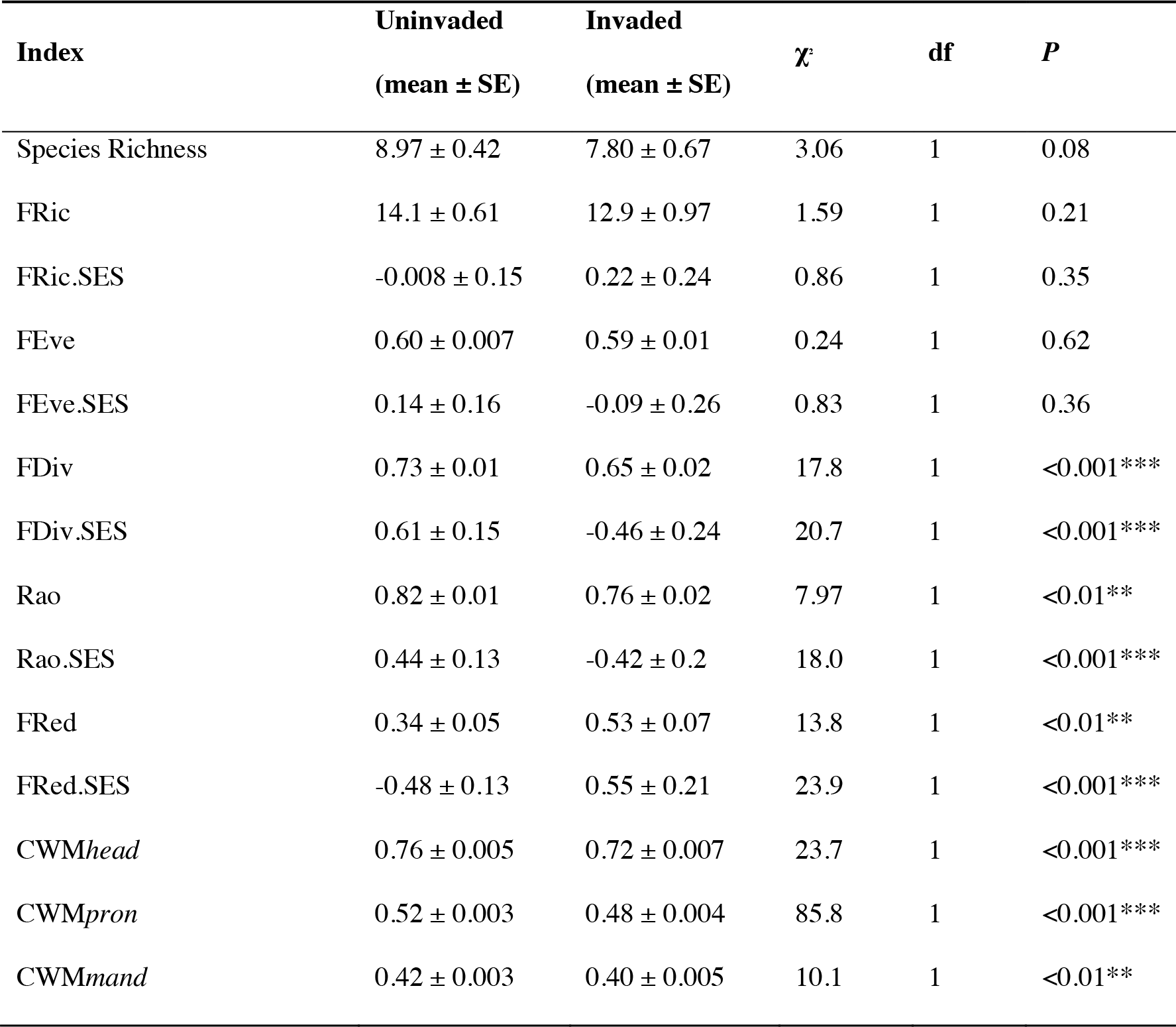
Summary statistics for response variables in separate linear mixed-effects models with community type (uninvaded vs. invaded) as fixed effects and environmental variation as a random effect.

**Figure 1.**
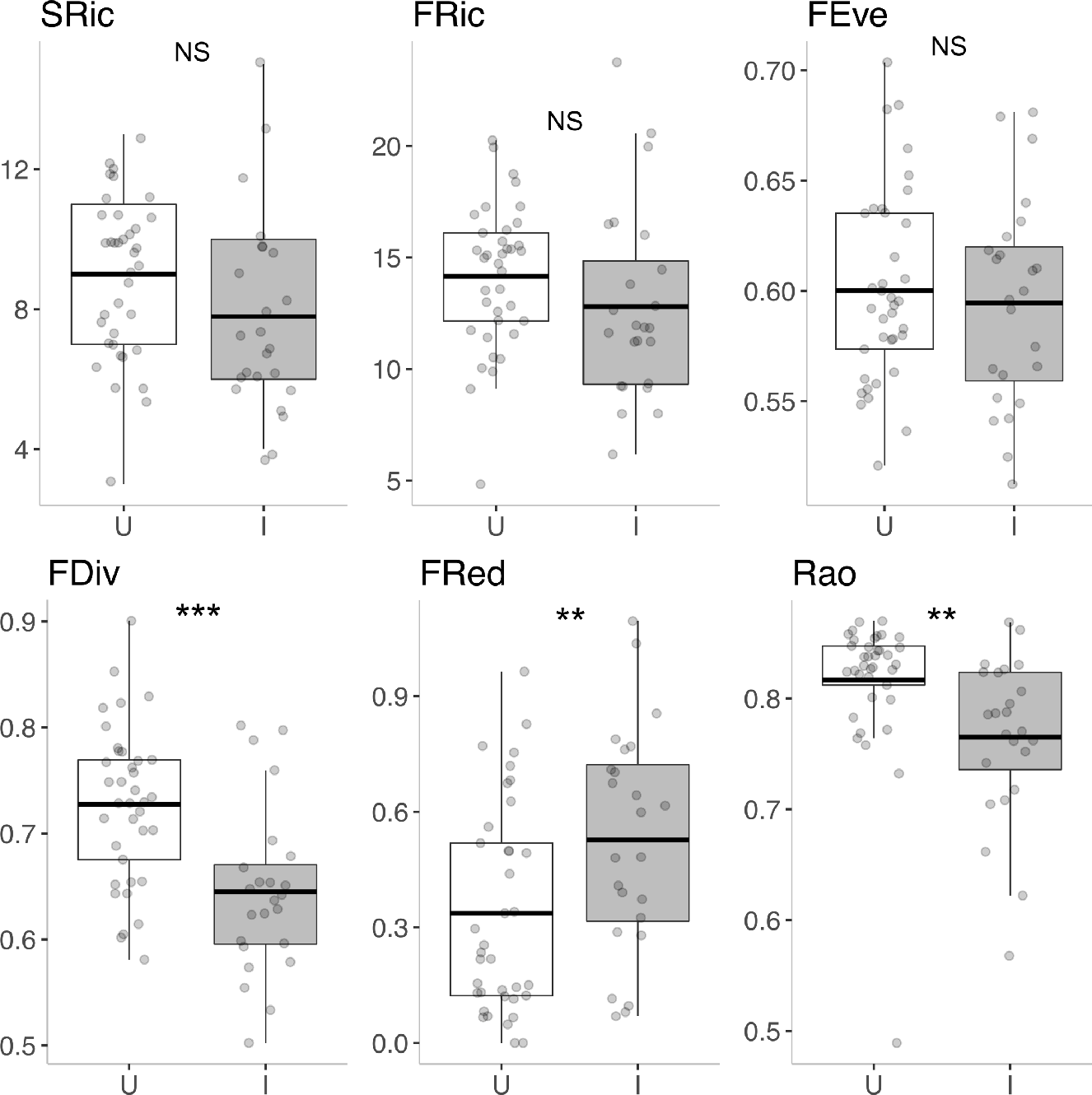
Boxplots showing species richness and observed values of five functional diversity indices in 37 uninvaded and 24 invaded local communities. Dots show values of individual communities, thick bars show means, boxes show inter-quartile range and vertical lines extend to maximum and minimum values (excluding outliers). Asterisks indicate statistical significance (*** *P*<0.001, ** *P*<0.01, * *P*<0.05, NS, not significant).

### Functional diversity and CWMs

Uninvaded and invaded communities had similar levels of FRic and FEve for both observed and SES values. However, in linear mixed-effects models, the observed FDiv and Rao of invaded communities were significantly lower than those of uninvaded communities by 11% and 7% respectively, and the FRed of invaded communities was significantly higher than that of uninvaded communities by 56% (Table 2; Fig. 1); similar relationships were observed for SES values. CWMs for the traits *size*, *scape length*, *eye width* and *leg length* did not differ significantly between uninvaded and invaded communities. By contrast, the CWMs for *head width*, *pronotum width* and *mandible length* were significantly smaller (by 4–7%) in the invaded communities (Table 2; Fig. 2).

**Figure 2.**
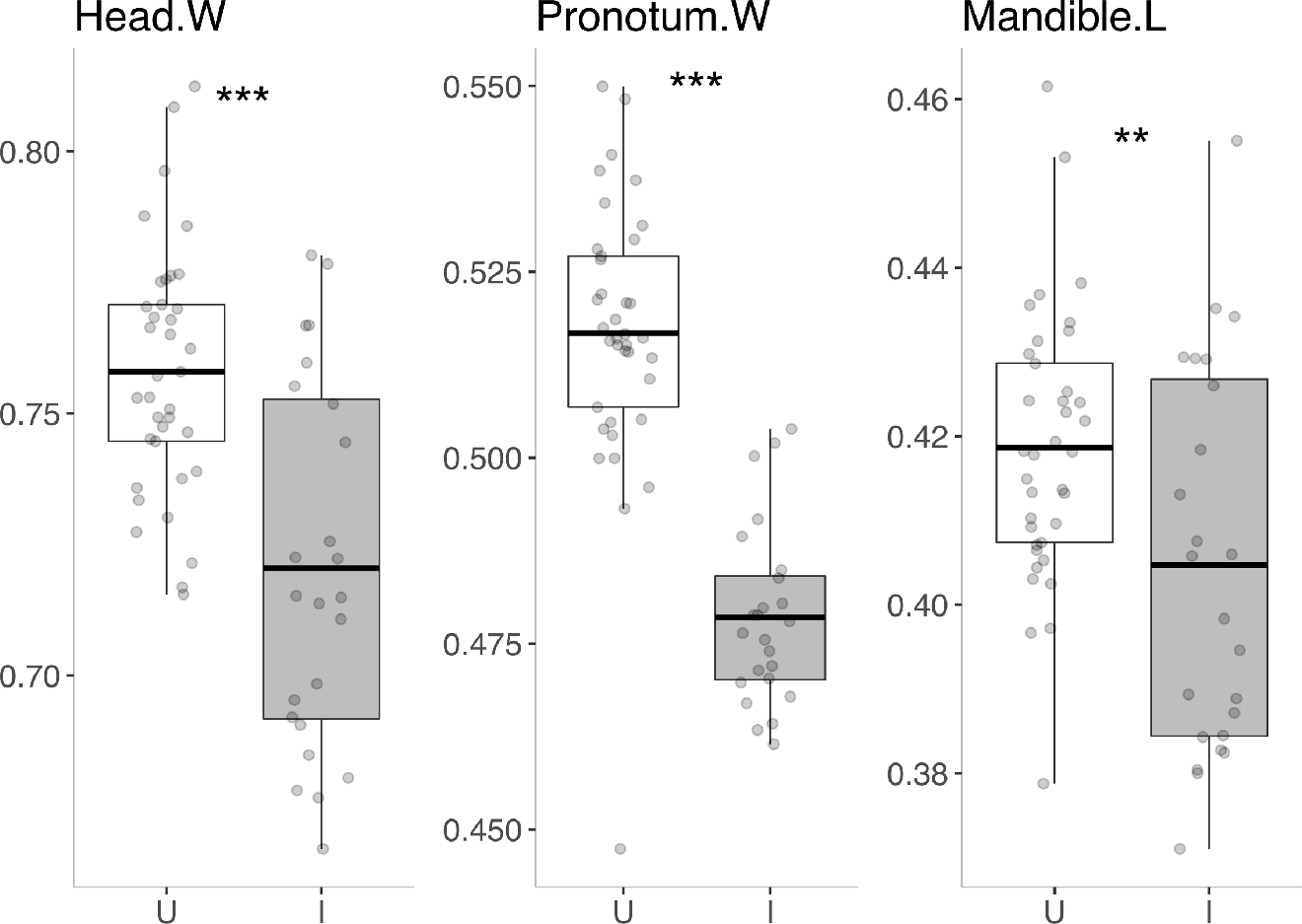
Boxplots displaying community-weighted mean values for size-corrected *head width*, *pronotum width* and *mandible length* in 37 uninvaded and 24 invaded communities. Dots show values of individual local communities, thick bars show mean values, box edges show standard deviations, and vertical lines extend towards minimum and maximum values. Asterisks indicate statistical significance (*** *P*<0.001, ** *P*<0.01).

### Taxonomic and functional beta diversity

Uninvaded and invaded communities were significantly dissimilar in both taxonomic and functional composition, and these dissimilarities were driven by turnover in species as well as trait values (Table 3; Fig. 3). The observed levels of total taxonomic and functional dissimilarities among both uninvaded and invaded communities were comparable (Table 4), but SES values revealed that total functional dissimilarity was lower among invaded communities when corrected for species richness (Mann-Whitney U test: *P* < 0.001) (Fig. 3). Invaded communities had significantly lower levels of functional turnover (by 23%) and higher functional nestedness (by 20%) in observed values; these relationships were maintained in SES values (Mann-Whitney U tests: *P* < 0.001) (Table 4). Likewise, invaded communities were significantly more taxonomically nested than uninvaded communities (by 42%, Table 4; Fig. 3). That is, in comparison to uninvaded communities, relatively greater proportions of the total taxonomic and functional dissimilarities among invaded communities were driven by losses of species than by replacements of species, and by changes in the abundances of trait values than by changes in the trait values themselves, respectively.

**Table 3.** Results of PERMANOVA tests for dissimilarities between uninvaded and invaded communities in their observed taxonomic and functional compositions.

| **Beta diversity** | **Component** | **F** | **R<sup>2</sup>** | **P** |
| --- | --- | --- | --- | --- |
| Taxonomic | Total Dissimilarity | 22.4 | 0.28 | <0.001*** |
|  | Turnover | 31.9 | 0.35 | <0.001*** |
|  | Nestedness | 13.6 | 0.31 | 1.0 |
| Functional | Total Dissimilarity | 33.1 | 0.36 | <0.001*** |
|  | Turnover | 32.7 | 0.36 | <0.001*** |
|  | Nestedness | 14.7 | 0.33 | 1.0 |

**Figure 3.**
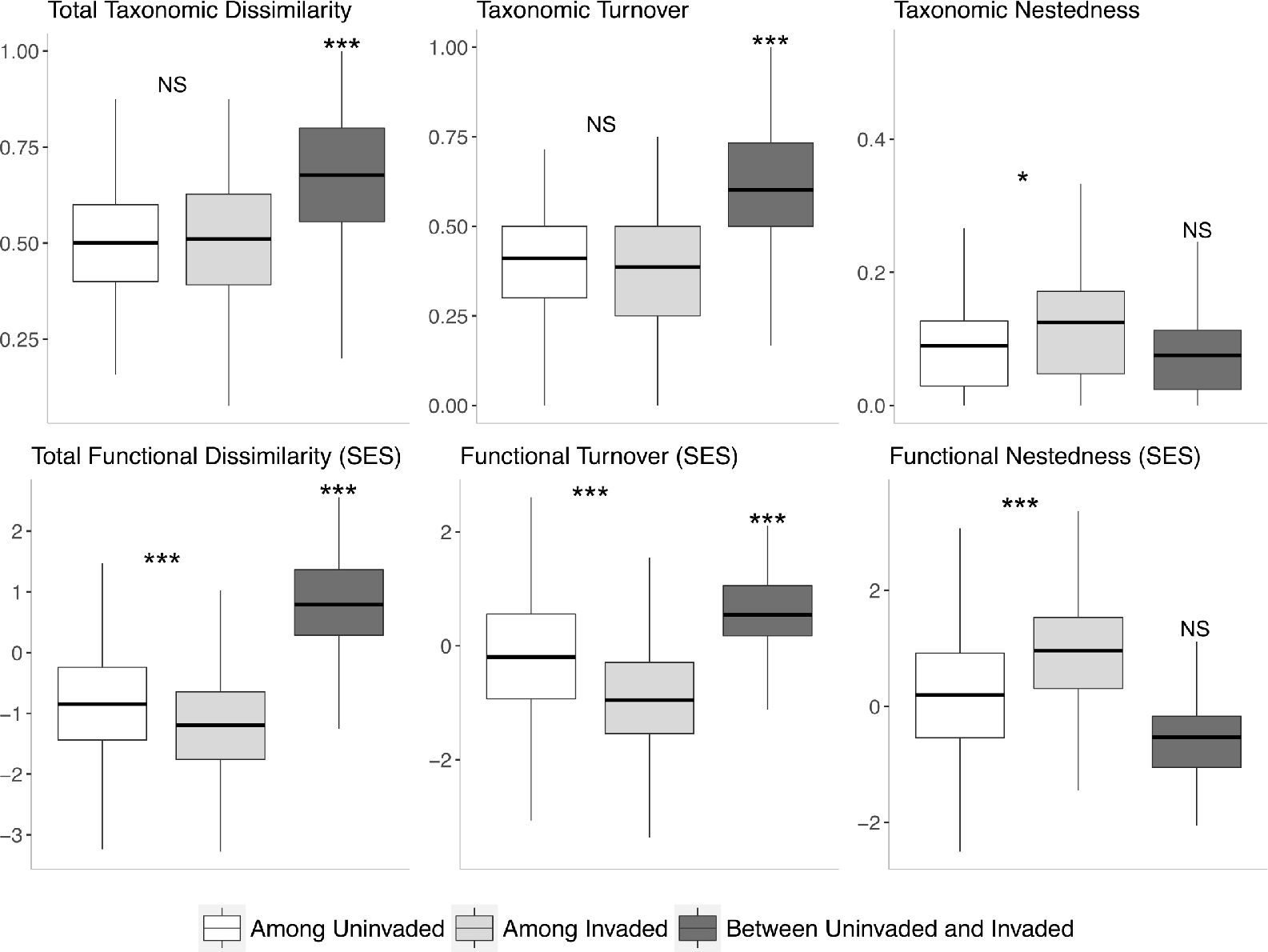
Observed levels of taxonomic beta diversity and functional beta diversity corrected for species-richness (SES values) in three measures of dissimilarity (Total, Turnover, and Nestedness). Boxplots show values among uninvaded communities, among invaded communities, and between uninvaded and invaded communities. Asterisks indicate statistical significance (*** *P*<0.001, * *P*<0.05, NS, not significant).

### Species’ functional richness and functional uniqueness

Functional richness varied over four-fold among species (Min. = 0.86, Max. = 3.61) (Fig. 4). The four most functionally-rich species were two dimorphic species of *Camponotus*, another dimorphic species, *Pheidole nodus*, followed by the polymorphic invader *S. invicta*. In separate linear regressions, species’ relative uniqueness to both uninvaded and invaded communities increased with their objective uniqueness (Fig. 5). However, there was relatively more overlap between the functional spaces of less unique species and the functional spaces of invaded communities (Intercept_Invaded_ = −0.15; Intercept_Uninvaded_ = 0.45). Furthermore, the relative uniqueness of species to invaded communities increased more steeply with an increase in objective uniqueness (Slope_Invaded_ = 1.17; Slope_Uninvaded_ = 0.52), such that very unique species were more unique to invaded communities than to uninvaded communities (Fig. 5).

**Figure 4.**
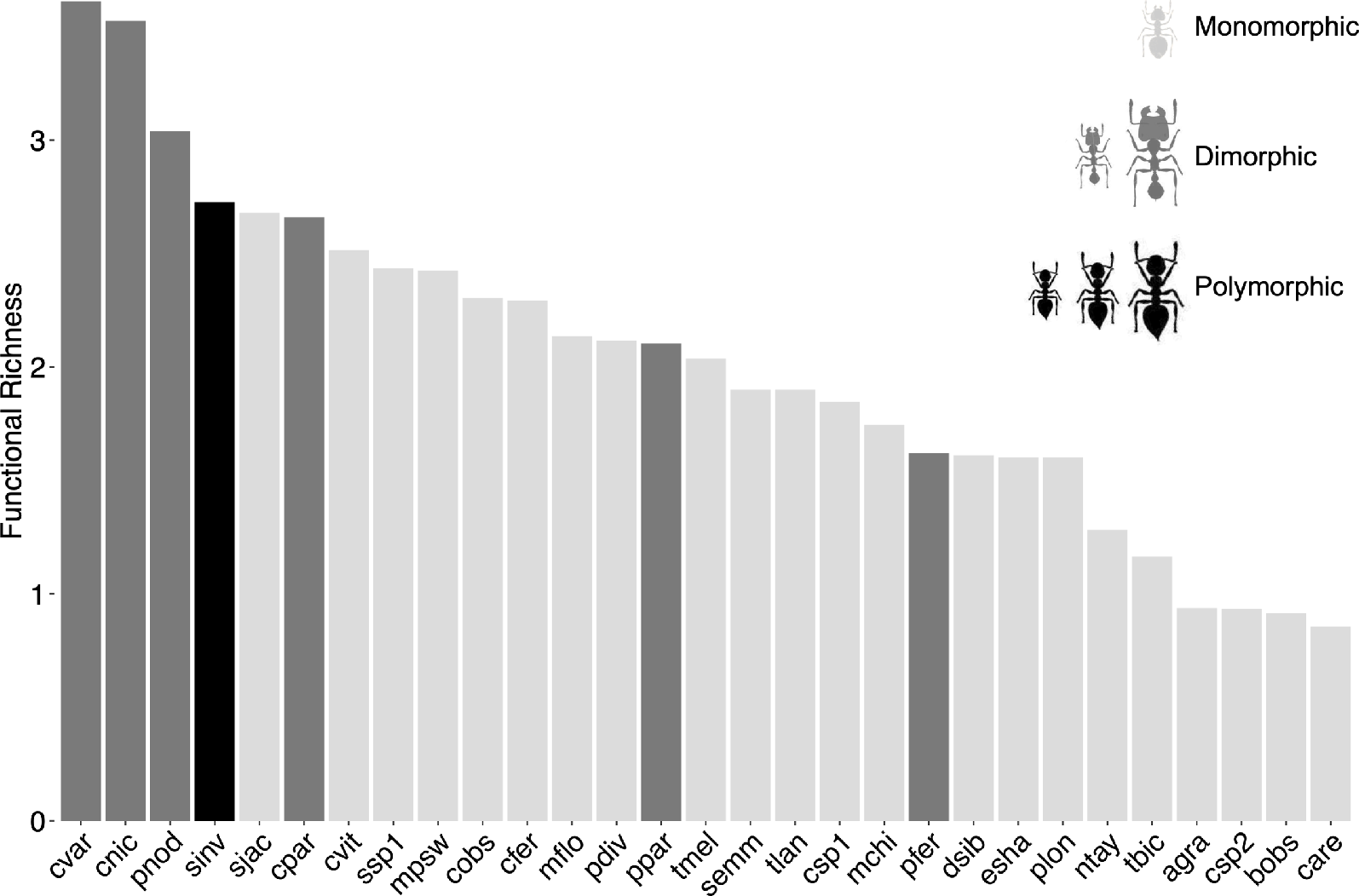
Functional richness of ant species with three different degrees of polymorphism. Bars show values for 22 monomorphic species (light grey), six dimorphic species (dark grey), and the polymorphic species *S. invicta* (black). Full species names are listed in Table S1, and images of monomorphic, dimorphic and polymorphic species are shown in Fig. S4.

**Figure 5.**
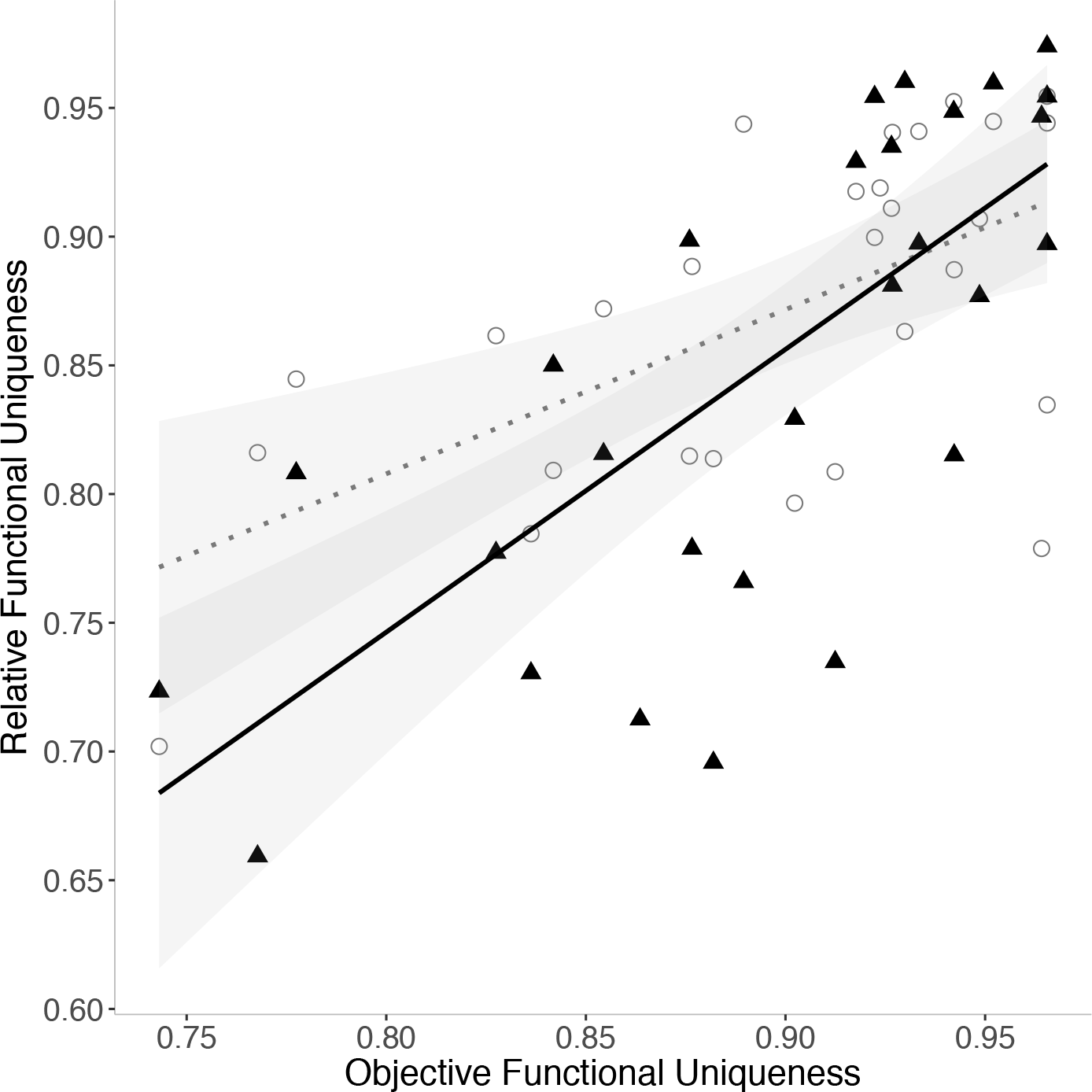
Species’ relative functional uniqueness to local communities that were uninvaded (grey dotted line; circles) and invaded (black solid line; triangles) plotted against their objective functional uniqueness in the species pool.

## Discussion

Invasions by alien species are recognized as top drivers of species extinctions and known to cause declines in the species richness of invaded communities (Bellard et al., 2016). However, invasions can also impact the structure and function of ecological communities in ways that are less detectable with taxonomic measures of diversity. Here we investigated the impacts of an invasion on the diversity and distribution of species and traits in individual communities (alpha diversity) and across multiple communities (beta diversity). We found that invasion by *S. invicta*, one of “the world’s worst invasive species” (Lowe et al., 2000), led to the functional homogenization and altered functional identity of ant communities in tropical Asia despite effecting marginal changes in species and functional richness. Additionally, we observed higher functional richness in polymorphic species than monomorphic species. Below we discuss the possible ecological drivers for the observed patterns and the general implications of our findings.

### Polymorphic species are more functionally rich

Polymorphisms are among the most conspicuous and pervasive sources of intraspecific trait variation (Ford, 1957). It is conceivable that polymorphic species, with greater variation in morphology, would display higher functional richness. We generally found this to be the case among the 29 ant species studied. The four most functionally-rich species were dimorphic species of the genera *Camponotus* and *Pheidole* and the polymorphic *S. invicta*, which displayed 2–322% more functional richness than monomorphic species (Fig. 4). However, the functional richness of three dimorphic species was higher than that of the polymorphic *S. invicta*. This likely resulted from the pronounced allometry of the discrete major subcastes of dimorphic species, which generated extreme trait values; polymorphic *S. invicta* workers, by contrast, display a continuous variation with size (Figs. 4 and S4). While it is unlikely that the morphological traits measured were representative of all species’ multidimensional niches, our findings demonstrate that, at least across multiple aspects of morphology linked to foraging, mobility and physiology (Table 1), polymorphic species occupy more functional space and may have access to a greater variety of niches than monomorphic species, as previously suggested (Farji-Brener et al., 2004). Future work investigating whether functional richness predicts niche variety or specialisation across polymorphic and monomorphic species could further our understanding of how species and ecosystem processes respond to ecological change.

### Lower species and functional richness in invaded communities

Disturbances drive local extinctions by decreasing the abundances of particular species with vulnerable trait combinations; this process can initially occur without significantly modifying community composition or affecting total species and functional richness (Mouillot et al., 2013). Invasions by *S. invicta* in Hong Kong generally follow these patterns. Two thirds of the native species had lower abundances in the invaded communities (Fig. S3), but the same communities displayed only marginally lower species richness (by 13%) and functional richness (by 8.5%) (Table 2 and Fig. 1). These results differ from previously reported extensive declines (by 69%) in the species richness of ant communities invaded by *S. invicta* in North America (Porter & Savignano, 1990). The current absence of an extensive decline in species richness may be associated with the relatively younger *S. invicta* invasion in Hong Kong (Ascunce et al., 2011), and potentially the ecological differences (and thus responses) between the tropical grassland ant communities studied here, which comprised many disturbance specialists, and the ant communities of temperate forests in previous studies (Porter & Savignano, 1990). Aside from the marginal differences in species and functional richness, multiple indices sensitive to changes in the abundance and distribution of trait values differed significantly between uninvaded and invaded communities (Table 2: CWMs, FDiv, Rao, and FRed). Although each index measures a unique aspect of functional identity and diversity (Mouchet et al., 2010), the results collectively indicate that invasions by *S. invicta* exert a non-random selection on native communities through the trait values of individuals; we discuss these patterns in the two sections that follow.

### Altered functional identity of invaded communities

The CWMs for size-corrected *head width*, *pronotum width* and *mandible length* of ants in invaded communities decreased significantly by 4–7% (Fig. 1), in line with our hypothesis that invasion would alter communities’ functional identities. The results suggest that invasion selects for individuals with narrower heads and pronotums and shorter mandibles. One hypothesis is that the observed patterns relate to mobility. It has been shown that the width of an ant’s head and pronotum determine the size of gaps through which it can pass (Schofield et al., 2016). Narrower heads and pronotums of ants in invaded communities could thus reflect demands for moving through tighter spaces to avoid the behaviourally dominant *S. invicta* during foraging (Tschinkel, 2006) or to reach resources in less accessible locations. For instance, native hypogaeic species, specialised to forage and move through soil, increased in relative abundance by 50% in the presence of other invasive ants (Human & Gordon, 1997). A second hypothesis relates to diet. In ants, long mandibles of many predatory species are specialized adaptations for prey capture (Silva & Brandão, 2010), and larger heads and pronotums afford more space for the musculature powering snapping, gripping and load-bearing abilities (Keller et al., 2014; Richter et al., 2019). Hence, the shorter mandibles and narrower heads and pronotums of ants in invaded communities may reflect a less specialized or more herbivorous diet. Such ants may use more liquid foods (e.g., honeydew from hemipterans) that can be ingested relatively quickly, and which require less manipulation (capture, ripping, transport) than solid foods. The presence of the behaviourally dominant *S. invicta* may select for ants that ‘grab and go’ over those which remain at food sources for longer periods.

### Functional clustering and redundancy of invaded communities

In invaded communities, values of FDiv and Rao decreased significantly by 11% and 7 % respectively, while FRed increased significantly by 56% (Fig. 2). These patterns arose due to the presence of more individuals with similar trait values in invaded communities. Specifically, FDiv reflects the degree to which the distribution of species’ abundances in functional space maximizes total community variation in trait values (and niches) (Mouchet et al., 2010). Hence, the results suggest that the most abundant species in uninvaded communities are more differentiated in their niches (higher FDiv), while those in invaded communities have more similar niches (lower FDiv). As observed for FDiv, the lower Rao in invaded communities is indicative of niche clustering (species’ abundances are relatively more clustered in functional space) (Mouchet et al., 2010). FRed reflects the degree to which specific trait values are represented by multiple species in the community (Carmona et al., 2016). In uninvaded communities (lower FRed) fewer species share the same trait values, suggesting less niche overlap. By contrast, more species share the same trait values in invaded communities (higher FRed), suggesting more niche overlap. Therefore, as observed for the CWMs, the results for FDiv, Rao and FRed show a general pattern of species’ abundances converging towards particular trait values in invaded communities, which is one signature of selection (Vellend, 2016).

### Functional homogenization across invaded communities

Our analysis of functional alpha diversity suggests that *S. invicta* invasions are associated with a selection for specific trait values in individual communities. Because such selection has repeated over separate communities invaded by *S. invicta*, functional beta diversity patterns across multiple communities show a trend towards functional homogenization. This is evident from the significantly lower functional dissimilarity among communities where *S. invicta* is present (Fig. 3). Contrary to our hypothesis, however, functional turnover did not track taxonomic turnover; changes in the species found in different invaded communities were not matched proportionately by changes in those communities’ trait values. The invaded communities actually retained similar levels of taxonomic turnover to uninvaded communities (Fig. 3). However, the former displayed significantly less functional turnover in observed structure (by 23%; Table 4), as well as in SES values of functional beta diversity corrected for the effects of species richness (Fig. 3). In previous analyses using computer simulations, such patterns of low functional turnover amid higher taxonomic turnover were predicted to emerge most frequently when there are high levels of functional redundancy in individual communities (Baiser & Lockwood, 2011). Given that communities invaded by *S. invicta* displayed 56% more functional redundancy than uninvaded communities (Fig. 2), our observations in an invasion context provide empirical support for the theoretical predictions (Baiser & Lockwood, 2011).

**Table 4.** Permutation tests for multivariate dispersions, with calculations based on the average distances to centroids of uninvaded and invaded communities for different components of taxonomic and functional beta diversity. These tests compare the levels of beta diversity observed among uninvaded communities to those observed among invaded communities.

| **Beta diversity** | **Component** | **Uninvaded** | **Invaded** | **F** | **P** |
| --- | --- | --- | --- | --- | --- |
| Taxonomic | Total Dissimilarity | 0.35 | 0.36 | 0.10 | 0.77 |
|  | Turnover | 0.29 | 0.28 | 0.20 | 0.66 |
|  | Nestedness | 0.07 | 0.10 | 5.89 | 0.02* |
| Functional | Total Dissimilarity | 0.32 | 0.31 | 0.01 | 0.91 |
|  | Turnover | 0.30 | 0.23 | 5.01 | 0.03* |
|  | Nestedness | 0.41 | 0.49 | 7.37 | <0.01** |

### Functionally ordinary winners and functionally unique losers across invaded communities

Invasions and other disturbances can result in losses of functionally unique species before functionally redundant species (Flynn et al., 2009). Thus, examining the abundance and distribution of functionally unique species may promote the advanced detection of invasion impacts. Previous studies used trait patterns of aggregated communities to define functionally unique groups or species, then analysed their abundances within each community (e.g., Coetzee & Chown, 2016). A species’ functional uniqueness, however, is a relative property, dependent on the value and abundance of other traits present in the particular community. Thus, also measuring species’ functional uniqueness as *relative to specific communities* may improve the understanding of changes in functional space and how shifting species abundances contribute to these changes. Here we first assigned each species an objective value of functional uniqueness in the species pool using a community containing all species at equal abundance. We then validated this measure by showing that objectively unique species were on the whole more unique than others across different uninvaded and invaded communities (Fig. 5: positive linear relationships for both lines). Next, we found that objectively non-unique (functionally ordinary) species constituted more of the functional spaces of invaded communities than uninvaded communities (Fig. 5: lower intercept of the ‘invaded’ line). We further found that objectively very-unique species constituted less of the functional spaces of invaded communities than uninvaded communities (Fig. 5: steeper slope of the ‘invaded’ line). Together, these findings suggest that the *S. invicta* invasion has led to invaded communities becoming more comprised of a subset of species (winners) sharing trait values which are common in the species pool, and less comprised of other species (losers) with trait values that are rare in the species pool. These patterns mirror the decline of functionally unique species before functionally redundant species observed in other disturbances (Flynn et al., 2009).

### Implications for ecosystem function

The consequences of an invasion will extend to the ecosystem if the affected taxa are also key modulators of ecosystem functions. Ants are such organisms, and the effects of *S. invicta* invasion on various ant-modulated ecosystem functions such as predation, nutrient cycling and bioturbation is a pertinent question to tackle in future research. These effects will hinge on the particular relationships between ant diversity and ant-modulated ecosystem functions in the tropical grassland communities studied here. For instance, if ecosystem functions mainly respond to the functional identities of the ant communities (i.e., selection effects), they may be impacted significantly by the altered CWMs of invaded communities. Alternatively, functional homogenization and the decline of functionally unique species in invaded communities could impact ecosystem functions driven by functional complementarity. Ecosystem functions may also respond to both functional identity and complementarity across different spatial and temporal scales (Isbell et al., 2018). Whatever the case, the present findings show *S. invicta* invasions in Hong Kong to have significant impacts on the functional structure of ant communities, which may also affect ecosystem-wide changes.

### Implications for mitigating the impacts of global species exchange

Using data on the traits of individuals we have shown that an alien invasive species alters – in a selective, non-random manner – the functional properties of native communities that may influence ecosystem processes. Crucially, our findings further indicate that such impacts may unfold in the absence of similar changes in both taxonomic and functional richness. Thus, assessments exclusively using taxonomic measures of diversity, or indices that only describe trait variety, may fail to detect various consequences of invasions for the structure and function of ecological communities. These other ecologically significant consequences of invasion (e.g., functional clustering and homogenization) can be uncovered by investigating patterns in the diversity and distribution of traits at the species, community, and landscape levels. As the global exchange of species continues to rise (Seebens et al., 2017), detecting harmful species early and evaluating their potential for impacts will only become more important. Using comprehensive trait-based approaches can help achieve these goals. This study demonstrates how trait-based approaches can improve understanding of the ways by which well-known invasive species like *S. invicta* impact the functional dimension of biodiversity. Furthermore, the results suggest that the same approaches may also be useful for identifying other potentially overlooked alien species driving cryptic impacts on ecosystem-relevant community structure amid modest changes to community metrics based on species’ taxonomic identities.

## Authors’ Contributions

M.K.L.W., B.G. and O.T.L. designed the study. M.K.L.W. conducted fieldwork, analysed the data and wrote the first draft of the manuscript. All authors contributed substantially to manuscript revisions.

## Acknowledgements

We are grateful to Brett Morgan for providing access to environmental data, Mac Pierce, Roger Lee and Roy Cheung for field assistance, Carlos Carmona for help with *TPD*, and staff of AEC and WWF Hong Kong for logistical support. This work was supported by a National Geographic Grant (60-16) and a University of Oxford Clarendon Scholarship to M.K.L.W.

